# An efficient and adaptable workflow for editing disease-relevant single nucleotide variants using CRISPR/Cas9

**DOI:** 10.1101/2021.11.12.467071

**Authors:** Inga Usher, Lorena Ligammari, Sara Ahrabi, Emily Hepburn, Calum Connolly, Gareth L. Bond, Adrienne M. Flanagan, Lucia Cottone

## Abstract

Single nucleotide variants are the commonest genetic alterations in the human genome. At least 60,000 have been reported to be associated with disease. The CRISPR/Cas9 system has transformed genetic research, making it possible to edit single nucleotides and study the function of genetic variants *in vitro*. While significant advances have improved the efficiency of CRISPR/Cas9, the editing of single nucleotides remains challenging. There are two major obstacles: low efficiency of accurate editing and the isolation of these cells from a pool of cells with other editing outcomes. We present data from 85 transfections of induced pluripotent stem cells and an immortalised cell line, comparing the effects of altering CRISPR/Cas9 design and experimental conditions on rates of single nucleotide substitution. We targeted variants in *TP53*, which predispose to several cancers, and in *TBXT* which is implicated in the pathogenesis of the bone cancer, chordoma. We describe a scalable and adaptable workflow for single nucleotide editing that incorporates contemporary techniques including Illumina MiSeq™ sequencing, TaqMan™ qPCR and digital droplet PCR for screening transfected cells as well as quality control steps to mitigate against common pitfalls. This workflow can be applied to CRISPR/Cas9 and other genome editing systems to maximise experimental efficiency.

**Simple Summary:** CRISPR/Cas9 has revolutionised genetic research. Cas9 generates a double strand break with high efficiency which is repaired by a cell’s pathways. If a genetic template is provided, the damage can be accurately repaired to introduce a desired genetic alteration. However, accurate repair occurs at a low efficiency and in a small proportion of edited cells, representing the main obstacles in harnessing CRISPR’s full potential. Using data from 85 CRISPR experiments for single nucleotide editing, targeting three locations in the human genome that are implicated in predisposition to cancer, we report the effect of different experimental conditions on editing efficiency. We describe current technologies that can be used to streamline the identification of accurately edited cells and synthesise these into an adaptable workflow that can be applied to CRISPR/Cas9 experiments to achieve single nucleotide editing in disease-relevant cell models.

## 1. Introduction

The Class 2 Clustered Regularly Interspaced Short Palindromic Repeat (CRISPR)/Cas9 system has transformed genetic research through a multitude of applications, from functional genomics to genome-wide screens and gene therapy. Despite exponential growth of the CRISPR field, leveraging CRISPR/Cas9 as a means of introducing a genetic alteration of interest with high efficiency and without off-target changes remains challenging [1].

The CRISPR/Cas9 system exploits the Streptococcus pyogenes-derived Cas9 endonuclease (SpCas9) enzyme to induce a double stranded break (DSB) at a specific genomic locus via a guide RNA (gRNA). A triplet of nucleotides, the protospacer adjacent motif (PAM), is required for the binding of Cas9 and cutting of DNA. The DSB, or cut site, is repaired by two main pathways: the dominant non-homologous end joining (NHEJ) pathway and the less efficient homology directed repair (HDR) pathway. NHEJ operates throughout the cell cycle, whereas HDR operates only during S and G2 phases, where the sister chromatid is available to be used as a template for accurate repair [2]. Therefore, repair of the DSB occurs predominantly via NHEJ whereas more faithful editing via HDR occurs as a rarer event. NHEJ ligates the cut ends of DNA to repair the DSB but is error-prone, introducing insertions and deletions (indels); this is employed to disrupt a gene’s sequence (knock-out) and silence its expression. In contrast, HDR repairs the damaged DNA accurately using a genetic template [3], a process that can be achieved *in vitro* by introducing an exogenous template, such as a single-stranded oligodeoxynucleotide (ssODN), also known as knock-in.

The efficiency of genome editing by CRISPR/Cas9 is highly variable, being affected by multiple factors [4,5] such as gene locus, nuclease and cell type [6]. The efficiency of NHEJ is up to 80% and is relatively feasible with current protocols. In contrast, successful editing only occurs at a rate of 2% to 5% for HDR and boosting this low efficiency remains an active area of research [4,5,7–15]. Experimental modifications that boost HDR rates fall into two categories: the design of CRISPR components (gRNA or ssODN) [4,5,9,10,12,14,15] and protocol adjuncts such as chemical inhibitors and temperature adjustments [7,8,13,16]. These modifications increase the rate of accurate repair (free of indels) by tipping the balance in favour of repair by HDR following a CRISPR/Cas9-induced DSB. Previous studies have focused on modifying the cut-to-mutation distance to increase knock-in efficiency, mostly utilising reporter assays to measure editing outcomes at the bulk population level without isolating viable clonal cell lines. Moreover, most studies have selected models, such as HEK293, known to be amenable to genetic manipulation, for optimisation purposes. Only a limited number of studies utilise other cells types such as induced pluripotent stem cells (iPSC) and embryonic stem cells, that are less susceptible to editing owing to their relative resistance to transfection, intact *TP53* pathway and intolerance of DNA damage [1,17].

Single nucleotide variants (SNVs) are the most common type of genetic sequence variation in the human genome. Over 60,000 SNVs are listed as pathogenic and are linked to human disease in the ClinVar database [18], but in many cases the role of such variants in the pathogenesis of the associated disease is unknown [19]. Modelling disease-associated variants *in vitro* represents a key experimental approach for elucidating their function [19] which provides valuable information for clinical management [20].

Many germline SNVs in *TP53*, the most commonly mutated gene in cancer, such as the hypermorphic G245D (rs121912656 SNV) and the R248Q (rs11540652 SNV) variants in exon 7, are recognised as being pathogenic and cause a range of cancers as part of the Li-Fraumeni syndrome [20]. Despite this, it is not known how these mutations bring about different cancer types. Modelling these variants in different cell types, and at different stages of differentiation, would allow the exploration of their tissue-specific effects. For many other SNVs, their association with disease has been established through large numbers of genome-wide association studies (GWAS) and other case-control studies, however their pathogenicity has not been robustly demonstrated. An example of such a SNV is the G177D (rs2305089 SNV) variant in exon 4 of *TBXT*, which is strongly associated with predisposition to the development of chordoma [21,22], a rare cancer of the spine. The mechanism by which this SNV is implicated in the pathogenesis of this disease is unknown.

The aim of this work was to improve HDR efficiency in CRISPR/Cas9 editing of single nucleotides. We focused on editing two genes of interest, the pathogenic alter G245D and R248Q in *TP53* and the likely pathogenic variant G177D in *TBXT*. To achieve this, we employed various contemporary technologies to screen mixed populations for the minor subset of edited cells and to ensure isolated cell lines are clonal and accurately edited.

## 2. Materials and Methods

### 2.1. U-CH1 chordoma cell line

#### Cell culture

The human U-CH1 chordoma cell line (ATCC® CRL-3217™ www.chordomafoundation.org) was grown as previously described [23]. Cell authentication was regularly performed by Short Tandem Repeat fingerprinting (Culture Collections, Public Health England, UK) (Supplementary Table 1).

#### CRISPR/Cas9 editing

All CRISPR/Cas9 components were purchased through Integrated DNA Technologies (IDT) (Coralville, IA, USA). ssODNs were ordered as Alt-R™ HDR Donor Oligos.

1. The gRNA was prepared by duplexing Alt-R® CRISPR-Cas9 crRNA and Alt-R® CRISPR-Cas9 tracrRNA (IDT, 1072532).
2. The ribonucleoprotein (RNP) complex was formed by combining 3.9μl Alt-R® CRISPR-Cas9 gRNA, 5.1μl Alt-R® S.p. Cas9 Nuclease V3 (IDT, 1081058) and 5.9μl sterile phosphate buffered saline for total volume of 15μl. All IDT components were resuspended in IDT duplex buffer according to the manufacturer’s instructions.
3. Upon reaching 80-90% confluence, cells were detached, counted, and transfected using the Lonza Amaxa® Cell Line Nucleofector® Kit V (Lonza, Basel, Switzerland VCA-1003) using electroporation program A30. To prepare the electroporation mixture a cell pellet of 2 million cells was resuspended in 70μl Lonza electroporation buffer, supplemented according to the manufacturer’s instructions, with 10μl of RNP complex and 3μl of modified Alt-R™ HDR Donor Oligo.
4. Following transfection, the transfected cells were recovered in medium without antibiotics.

#### Analysis of editing outcomes using digital droplet polymerase chain reaction (ddPCR)

A common primer set and probes for each allele were designed: a hexachlorofluorescein (HEX) probe for the parental allele (A/T) and fluorescein amidite (FAM) probe for the edited allele (G/C). All ddPCR assays were designed using primer3plus [24] according to the criteria and settings on pages 11 to 13 of the BioRad Droplet Digital™ PCR Applications Guide (http://www.biorad.com/webroot/web/pdf/lsr/literature/Bulletin_6407.pdf). All ddPCR experiments were carried out using the BioRad QX200 workflow employing the

Automated Droplet Generator using BioRad Automated Droplet Generation Oil for Probes (BioRad, Hercules, California, USA; #1864110), Eppendorf vapo.protect thermocycler and QX200 Automated Droplet Reader. The results were analysed using the BioRad QuantaSoft™ Analysis Pro Software using the rare event detection setting. The ddPCR supermix for probes (no dUTP) workflow was used.

#### Flow cytometry Activated Cell Sorting (FACS)

After transfection, U-CH1 cells were recovered for 24 to 36 hours before being single cell sorted into collagen-coated 96 well plates using a BD FACS Aria Fusion Cell Sorter™ (Becton Dickinson, Franklin Lakes, New Jersey, USA) running FACSDiva Software version 6. Cells were sorted to exclude TOPRO3+ dead cells and to select the top 10% of ATTO-550 positive cells that contain the IDT RNP complex.

#### Mirror plate for DNA extraction

Cells were washed with 100μl DPBS then incubated in 50μl of Accutase® (Innovative Cell Technologies, Inc., San Diego, CA AT104) at 37°C for 10-15 minutes, after which 50μl of medium was added. 50μl were taken for genomic DNA extraction using Lucigen QuickExtract™ (LGC, Middlesex, UK, QE09050) while the other 50μl were placed into a collagen-coated 96 well plate for subculture. 50μl of Lucigen QuickExtract™ can be added directly to the medium containing detached cells for rapid DNA extraction.

### 2.2. Induced pluripotent stem cells (iPSC)

#### Cell culture

The human episomal line of induced pluripotent stem cells (A18945, Gibco; Thermo Fisher Scientific, Inc., Waltham, MA, USA) was maintained in feeder-free culture on Geltrex matrix (A1413202, Invitrogen, Thermo Fisher Scientific) and Essential 8 Flex (E8 Flex) medium (A28585, Gibco, Thermo Fisher Scientific) supplemented with 0.5% of Penicillin (10,000 U/ml) unless otherwise noted. iPSCs used in this study were between passages 40 and 75. Cells were cultured at 37°C in a humidified atmosphere with 5% CO2. Upon reaching 80-90% confluence, usually within 3 to 4 days of culture, before colonies overgrew or began to differentiate in the centre, cells were passaged with a split ratio ranging from 1:3 to 1:6, by incubation for 5 minutes at 37°C with DPBS-EDTA 0.5mM pH 8.00 (Invitrogen, Thermo Fisher Scientific, 14190250 and 15575020).

#### CRISPR/Cas9 editing

All CRISPR/Cas9 components were purchased through IDT. ssODNs were ordered as Alt-R™ HDR Donor Oligos.

1. The gRNA was prepared by duplexing Alt-R® CRISPR-Cas9 crRNA and Alt-R® CRISPR-Cas9 tracrRNA (IDT, 1072532).
2. The RNP was formed by combining 0.78μl Alt-R® CRISPR-Cas9 gRNA, 1.02μl Alt-R® S.p. Cas9 Nuclease V3 (IDT, 1081058) and 1.2μl sterile phosphate buffered saline for total volume of 3μl. All IDT components were resuspended in IDT duplex buffer according to the manufacturer’s instructions.
3. Upon reaching 80-90% confluence, cells were detached, counted and transfected using the Lonza™ P3 Primary Cell 4D-Nucleofector™ X (Lonza, V4XP-3024) and electroporation program CA137.
4. To prepare the electroporation mixture, a cell pellet of 0.5 million cells was resuspended in 20μl Lonza electroporation buffer, supplemented according to the manufacturer’s instructions, 1μl of RNP complex and 0.5μl of the Alt-R™ HDR Donor Oligo.
5. Following transfection, cells were recovered in the following conditions: (i) 37°C, (ii) 32°C for 24 hours then returned to 37°C, (ii) in the presence of Alt-R® CRISPR-Cas9 HDR enhancer (IDT, 1081072) at the recommended concentration of 30μM in E8 Flex with RevitaCell™ but without antibiotics for 24 hours, (iv) DMSO at 1% in E8 Flex RevitaCell™ but without antibiotics for 12 to 24 hours. 12 to 24 hours after electroporation the media was changed to E8 Flex without HDR enhancer or DMSO.

#### Colony picking

Cells were detached with Accutase® (Innovative Cell Technologies, AT104), collected in E8 Flex plus RevitaCell™, dissociated into single cells, counted and plated at low density (700-1000 cells per dish) in 10-cm Geltrex-coated dishes, then maintained in fresh E8 Flex medium with RevitaCell™. When colonies appeared, usually around 4 days after plating, the medium was changed to E8 Flex without RevitaCell™ until the colonies were large enough to be picked. After 10 to 12 days, the medium was changed to E8 Flex plus RevitaCell™ for 2-3 hours before medium-sized colonies with reasonable morphology were picked under direct vision using a P200 pipette. The aspirated colony was transferred to a 96 well plate, containing 100μl of E8 Flex with RevitaCell™, and triturated up and down 10 times. The day after, medium was replaced by E8 Flex without RevitaCell™ and every other day until cells were 50-70% confluent and ready to be split 1:2. Half were seeded into a 96 well mirror plate for genomic DNA extraction and half for subculture or freezing.

#### Mirror plates for freezing and for DNA extraction

Colonies expanded in 96 well plates were pre-treated with E8 Flex with RevitaCell™ for 2-3 hours, washed gently with 100μl DPBS, and incubated in 30μl of Accutase® at 37°C for 10-15 minutes. A mirror plate for freezing was prepared containing 50μl of 2X freezing medium (E8 Flex plus 20% DMSO) and kept on ice. Cells were collected with 70μl of E8 Flex with RevitaCell™ by pipetting up and down and 50μl were transferred to the mirror plate and stored in Styrofoam container at −80°C. The remaining 50μl of suspension was kept in the original plate for genomic DNA extraction: the plate was spun at 1950 RCF for 30 minutes at 4°C, the medium was removed, and the plate was placed at −80°C. Upon thawing 30-50μl of Lucigen QuickExtract™ DNA Extraction Solution was added on ice to each well, triturated and extracted as per the Lucigen protocol below. The extracted DNA was then directly used for PCR or stored at −80°C.

### 2.3. Techniques common to both cell models

Regular testing was performed to ensure that the U-CH1 and iPSC lines were free of mycoplasma contamination using the EZ-PCR Mycoplasma Test Kit (K1-0210, Geneflow, Lichfield, Staffordshire, UK).

### 2.4. Genotyping using Illumina MiSeq™ next generation sequencing (NGS)

DNA was extracted using Zymo Column Extraction (Zymo Research, Irvine, California, USA, D3024) according to manufacturer’s instructions (for bulk transfections) or using the Lucigen QuickExtract™ DNA Extraction Solution (for picked colonies). PCR was performed with Kapa Hifi HotStart polymerase using conditions recommended for a <500 base pair (bp) product (Kapa Biosystems, Roche Molecular Systems, Inc., Pleasanton, California, USA, KR0370) using the following: 12.5μl 2X KAPA HiFi HotStart ReadyMix, 0.75μl 10μM Forward Primer (with MiSeq™ adapter, Supplementary Table 1), 0.75μl 10μM Reverse Primer (with MiSeq™ adapter), 2μl DNA and PCR-grade water up to 25μl. 3μl of DNA extracted in Lucigen was used for PCR. The PCR products were purified using the QIAquick PCR Purification Kit according to manufacturer’s instructions (QIAGEN Ltd., Manchester, England). MiSeq™ was performed in-house as described in [25].

### 2.5. Analysis of editing outcomes using MiSeq™ data

Paired-end FASTQ files were analysed using Cas Analyser [26] (http://www.rgenome.net/cas-analyzer/#!) with the following parameters: Nuclease type= single nuclease, comparison range (R) = 40, Minimum frequency (n) =1 and no optional wild type marker. Rates of unedited and edited outcomes (NHEJ +/− substitution and HDR) were calculated by number of reads containing outcome / total number of reads. Results are shown as percentages.

Unedited reads are defined as those which match the reference sequence in composition and length of sequence. NHEJ reads are defined as those which show insertions and/or deletions (change in sequence length compared to reference) but no knock-in of the desired mutation. Combined reads are those that contain indels and the desired mutation (also known as mixed repair reads, see [5]). HDR reads are those that contain the desired mutation and no off-target alterations or sequence length changes.

For editing outcomes of the colony-picked iPSC clones we defined the outcomes as follows:

- Homozygous unedited (90 to 100% of reads match the reference sequence)
- Homozygous knock-out (90 to 100% of reads show indels)
- Heterozygous unedited and knock-out (40 to 60% of the reads match the reference sequence and 40 to 60% show indels)
- Heterozygous knock-in (40 to 60% of the reads match the reference and 40 to 60% show the knock-in)
- Heterozygous knock-out (40 to 60% of the reads match the reference sequence and 40 to 60% show indels)
- “Hemizygous” (40 to 60% of the reads show indels and 40 to 60% show knock-in)
- Mixed repair (more than or equal to 40% reads in each category)

### 2.6. Genotyping using Sanger sequencing

PCR using the DNA extracted with Lucigen QuickExtract™ was performed as follows: 12.5μl AmpliTaq Gold™ 360 Master Mix (Thermo Fisher Scientific, 4398881), 0.5μl 10μM forward-reverse primer mix (Supplementary Table 2) and 10μl water plus 2μl DNA. PCR products were cleaned up using the ExoSAP-IT™ Express PCR Product Cleanup Reagent (Applied Biosystems, Thermo Fisher Scientific, 15563677) according to the manufacturer’s instructions and sent for Sanger sequencing (Source BioScience, Nottingham, United Kingdom).

### 2.7. Genotyping using TaqManTM qPCR

DNA was extracted as above and TaqMan™ genotyping was performed for rs2305089 (Applied Biosystems, 4351379) as follows: 5μl TaqMan™ genotyping mastermix, 0.5μl Taqman™ primer/probe mix (C__11223433_10), 3.5μl water, 1μl DNA. Results were interpreted using the Genotyping application on the Thermo Fisher Connect™ cloud.

### 2.8. Data analysis

All analysis and statistics were performed using R version 4.0.5 (2021-03-31). Pie charts were created with GraphPad Prism version 8.0.0 for Windows (GraphPad Software, San Diego, California USA, www.graphpad.com). All cartoons were created with Biorender.com.

## 3. Results

### 3.1. Editing of *TP53* SNVs in iPSC

We tested the ability of various CRISPR/Cas9 strategies to introduce the pathogenic G245D and R248Q germline variants into iPSC (Figure 1a). iPSCs were chosen because they express high levels of TP53, which correlate with the presence of open chromatin at this gene locus thereby allowing access of the CRISPR/Cas9 complex. Moreover iPSCs can be differentiated into a range of cell types, allowing the study of these variants in tissue-specific contexts, reflecting their role in the predisposition to a range of cancers [27].

**Figure 1.**
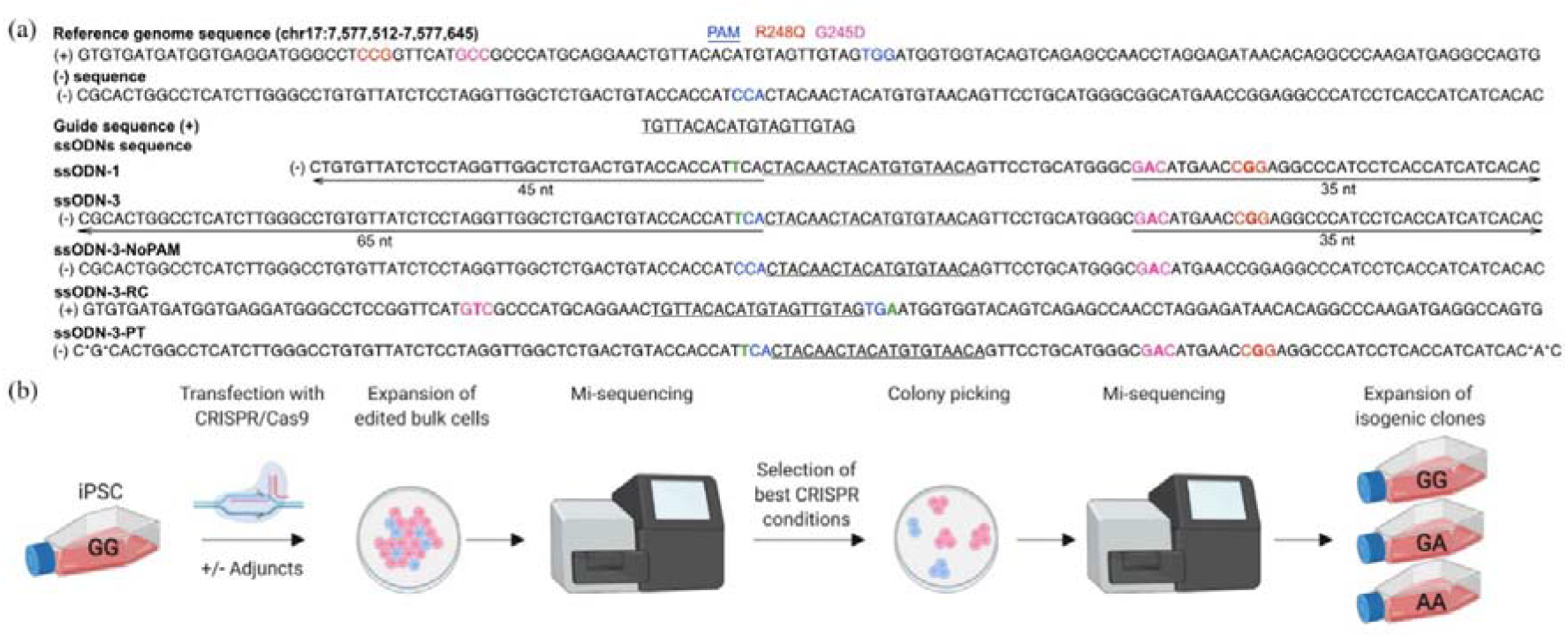
Workflow for early characterisation of editing outcomes followed by isogenic subline isolation in iPSCs. (a) Design of CRISPR/Cas9 components. Five ssODN designs were compared. *: phosphorothioate bonds. (b) iPSCs were transfected with the ssODNs under different experimental conditions. Bulk populations were screened by MiSeq™. Populations with the highest rates of HDR were selected for colony picking, which were then screened using MiSeq™. Individual colonies showing accurate repair were expanded as isogenic lines.

Recognising that iPSCs accumulate genetic alterations during cell culture [28,29], including SNVs at G245D, R248Q and other loci in *TP53* [29], we first ensured that a genetically pure population of iPSC was utilised for transfection. This was achieved by picking 20 clonal sublines, grown from the parental iPSC line; five sublines were screened by Sanger sequencing, all of which were free of SNVs in the 300bp surrounding the G245D and R248Q loci. One of these five sublines was selected for further experiments (Supplementary Figure 1).

### 3.2. Design of CRISPR/Cas9 gRNA and ssODN in iPSCs

We employed the IDT RNP system (Alt-R™ CRISPR-Cas9 System), composed of a gRNA that is composed of a CRISPR RNA (crRNA) hybridised to the 67mer tracrRNA as a duplex, which activates the Cas9 protein. The universal tracrRNA contains the ATTO™ 550 fluorophore which was used to check transfection efficiency by microscopy and by flow cytometry: the transfection efficiency approached 100% in iPSCs by flow cytometry (Supplementary Figure 2a).

We designed six gRNAs to target the region containing G245D and R248Q in *TP53* using E-CRISP (http://www.e-crisp.org/E-CRISP/) [30], CHOPCHOP (https://chopchop.cbu.uib.no/) [31] and the IDT Alt-R™ CRISPR HDR Design Tool (https://eu.idtdna.com/pages/tools/alt-r-crispr-hdr-design-tool) (Supplementary Figure 2b). Candidate gRNAs were found to be comparable when assessed *in silico* for the likelihood of on- and off-target effects using the IDT CRISPR-Cas9 guide RNA design checker (https://eu.idtdna.com/site/order/designtool/index/CRISPR_SEQUENCE). We then tested the six candidate gRNAs *in vitro*: we transfected RNPs, containing each of the six gRNAs individually, followed by Sanger sequencing. Only one gRNA was successful in producing a DSB as demonstrated by evidence of indels at the desired location (Supplementary Figure 2c and Figure 1a).

Previous studies have demonstrated that increasing distance from the target mutation to the cut site (cut-to-mutation distance) reduces efficiency of accurate HDR [2,4,5,10]. Studies that reported high HDR efficiency introduced mutations in close proximity (<5 nucleotides) to the cut site [7,16]. However, in our experiments, while it was possible to design guides that were predicted *in silico* to cause cleavage of the DNA close to the G245D or R248Q SNVs, the only gRNA that caused a DSB *in vitro*, did so at a distance of >15 nucleotides from the SNVs (Figure 1a and Supplementary Figure 2b).

### 3.3. Optimising the efficiency of accurate HDR in iPSCs

As we could not modify the cut-to-mutation distance, we tested if modification of the ssODN design or experimental conditions could improve the rate of HDR. We performed 75 individual transfections of iPSCs using electroporation (Figure 1a-b) (Table 1). Using Illumina MiSeq™ next generation sequencing (NGS) we analysed the results at the level of the bulk population with dedicated computational tools (for a review of tools see [32]). Transfected cells were frozen while awaiting MiSeq™ results.

**Table 1.** Modifications to the CRISPR/Cas9 protocol to boost editing rates.

| Modification | Mechanism(s) of action |
| --- | --- |
| gRNA design [15] | Preferential binding of CRISPR/Cas9 |
| Asymmetric ssODN [2,14] | Influences annealing and release of strands being repaired |
| Blocking mutation in ssODN [5,12] | Prevents cleavage of ssODN by Cas9 |
| Phosphorothioate modification of nucleotides [9,33] | Prevent degradation of ssODN |
| ssODN complementary to non-target strand [14] | Preferential binding of CRISPR/Cas9 |
| Cold shock [8,11] | Not known, possibly accumulation of cells in G2/M phase or persistence of RNP |
| HDR enhancer [11,13], reviewed in [34] | Inhibits NHEJ |
| Changing length of homology arms [12] | Influences annealing and release of strands being repaired |
| DMSO [35] | Not known, possibly improved DNA access or cell cycle arrest |

The editing efficiency across all ssODN designs and experimental conditions was variable, ranging from 0% to 12% for accurate HDR (Figure 2a-b). Comparing ssODN designs, we corroborated previous reports [10,14] that asymmetry of the homology arms is associated with superior rates of accurate HDR (range 1-12%, mean 5.2%, 40 samples) compared to symmetric design (range 0-6%, mean 3.5%, 11 samples) (Figure 2a). We then explored whether additional modifications to the asymmetric ssODN would increase the rate of accurate HDR further. These were the introduction of a silent mutation of the PAM [12], addition of phosphorothioated nucleotides [9,33] or the reverse complement (RC) of the asymmetric design [14] (Figure 1a, Table 1 and Supplementary Table 2). None of these modifications were taken forward as they did not improve the HDR rate (Figure 2a).

**Figure 2.**
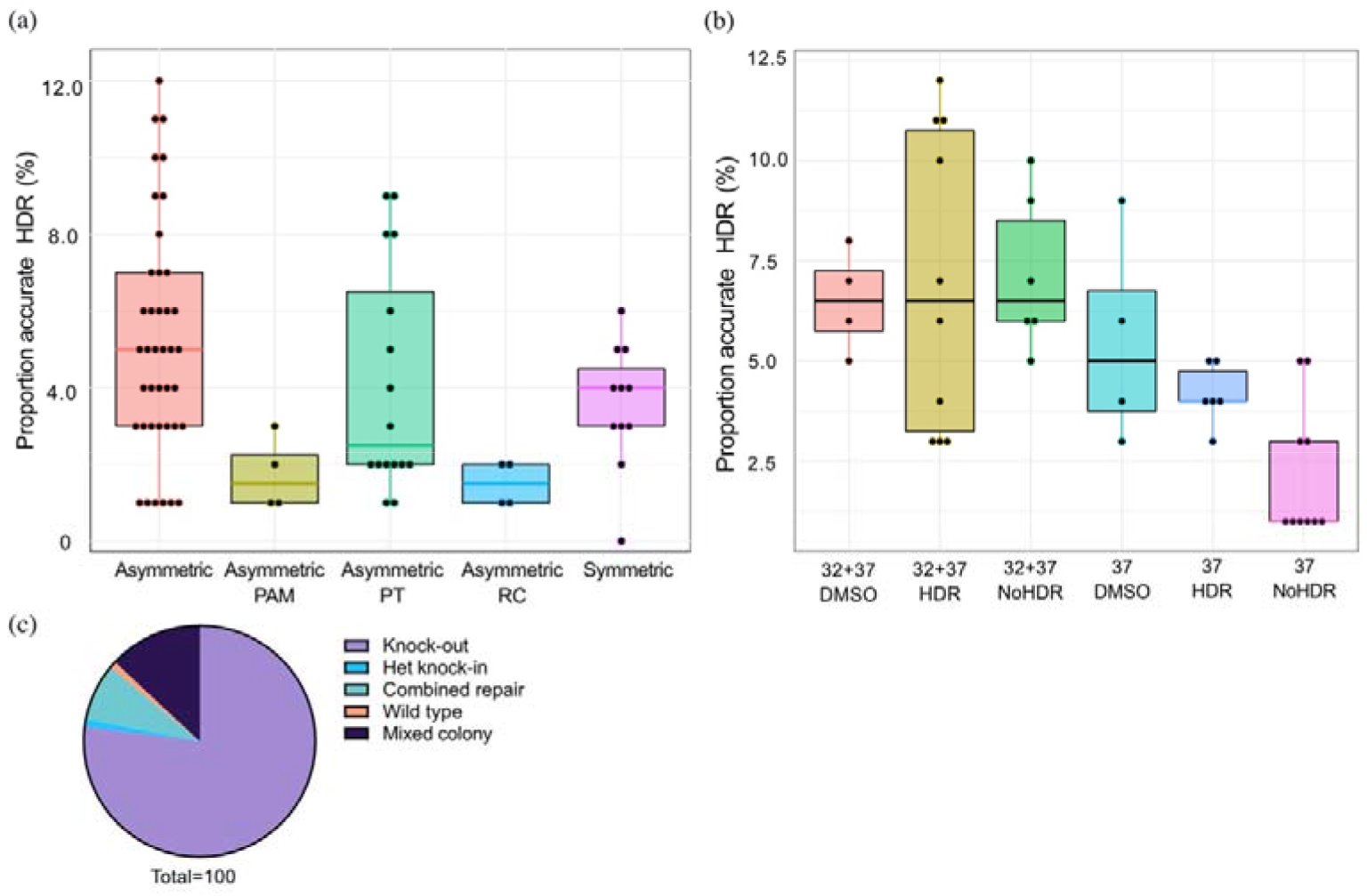
Comparison of HDR efficiency associated with protocol modifications at the bulk population level and editing outcomes in clonal lines. (a-b) Proportion of reads showing accurate repair by HDR in bulk population DNA transfected with (a) different ssODNs and (b) different experimental conditions. Asymmetric PAM: asymmetric donor without blocking mutation in PAM. Asymmetric PT: addition of phosphorothioate nucleotides to asymmetric ssODN. Asymmetric RC: reverse complement of asymmetric ssODN. 32+37: cold shock. HDR: addition of Alt-R™ HDR Enhancer after transfection. DMSO: addition of DMSO after transfection. NoHDR: no HDR enhancer or DMSO after transfection. (c) Editing outcomes in 100 colonies picked from two transfections showing HDR rates of 11 and 12%.

Next, using only the asymmetric ssODN, we proceeded to modify post-transfection experimental conditions by adding 1% dimethyl sulfoxide (DMSO) or the IDT Alt-R™ HDR Enhancer [35] and culturing cells at 32°C for 24 hours (cold shock) [8] (Table 1). We found that the most effective protocol included both cold shock and treatment with Alt-R™ HDR Enhancer, a finding consistent with previous studies [7,16] (Figure 2b). We also confirmed reports by others that DMSO was an effective chemical adjunct for increasing HDR rates, making it a cost-effective alternative to commercial HDR enhancers if tolerated by cells [35] (Figure 2b).

### 3.4. High efficiency of knock-in together with indels (combined repair process) in iPSC clones

We next proceeded to isolate isogenic cell lines from the bulk population to determine if the HDR efficiency achieved in the bulk cell population was translated to viable clonal lines which would be taken forward for experiments. Notably, most previously published protocols do not report details of clonal line isolation. We chose two transfected bulk populations which showed a promising rate of accurate HDR (11 and 12%) for single cell line isolation. These cells were thawed, plated at low density and 100 colonies were manually picked and expanded in 96 well plates then screened by MiSeq™ (Figure 1b and Figure 2c). We found a high rate of repair by NHEJ (77/100 clones, 77%). 13/100 (13%) colonies instead showed sequencing reads suggestive of a mixed population (see Methods), which if desired, could be subjected to further colony picking to purify the outcomes. A combined repair process with incorporation of the single nucleotide edit in addition to repair by NHEJ, was found in 8/100 (8%) colonies. One colony of 100 showed no sign of editing (wild type) and one colony was found to harbour a heterozygous knock-in free of indels: both colonies were generated by a protocol with an asymmetric donor, HDR enhancer and cold shock. The efficiency of accurate HDR was 1%.

Finally, quality assurance of the clonal lines was undertaken to assess the homogeneity/purity of the population and identify off-target alterations. Using MiSeq™ data we established that the wild type and the heterozygous iPSC clonal lines were genetically pure and free of off-target alterations in the 200 bp surrounding the variants.

In summary our data show that rates of HDR vary between transfections but can be boosted with asymmetric donors and HDR-enhancing modifications. Screening by MiSeq™ is useful for identifying populations with the highest rates of HDR before proceeding to isolating clonal lines as there is an attrition in the rate of HDR from bulk to viable cell line isolation.

### 3.5. Editing of *TBXT* in the U-CH1 immortalised cancer cell line

As the G177D SNV in *TBXT* is associated with a single cancer, chordoma, we employed the U-CH1 chordoma cell line which expresses TBXT at high levels and is largely faithful to the tumour genomically [32]. The U-CH1 cell line has two copies of *TBXT* which maximises the probability of successful editing compared to cell lines harbouring multiple copies.

### 3.6. Design of CRISPR/Cas9 for the U-CH1 chordoma cell line

We employed the IDT RNP system (Alt-R™ CRISPR-Cas9 System) and undertook 10 transfections by electroporation, similar to that used for iPSCs (see above). Prior to starting this workflow, we first ensured a pure population with respect to the region surrounding the G177D variant using MiSeq™ (Supplementary Figure 3a).

Four candidate gRNAs were suggested by the same computational tools used for iPSCs and were assessed for their on- and off-target scores by IDT CRISPR-Cas9 guide RNA design checker. The gRNA predicted to have the highest on-target score *in silico* was the only one that caused a DSB when tested *in vitro* (Figure 3a and Supplementary Figure 3b-c). As introducing a silent mutation into the PAM in the ssODN was not possible without altering the coding sequence, we modified another nucleotide in the ssODN to introduce a silent mutation which would still be expected to prevent binding and re-cleavage by the CRISPR/Cas9 complex (Figure 3a) [4,5,12].

**Figure 3.**
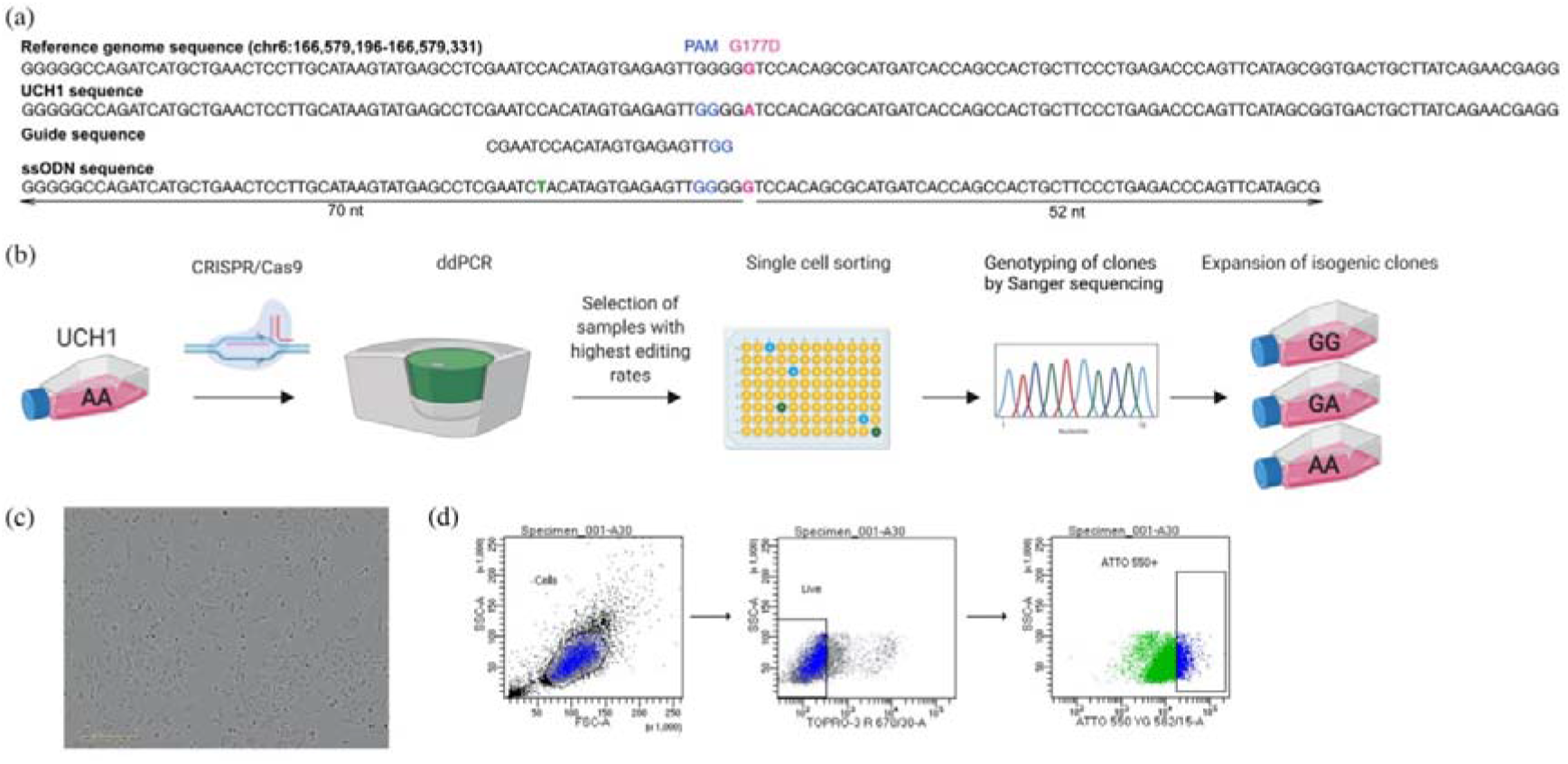
Workflow for early single cell sorting and high throughput genotyping. (a) Design of CRISPR/Cas9 components targeting the G177D SNV in *TBXT*. The silent mutation introduced in the ssODN is shown in green. (b) After transfection, U-CH1 bulk populations were screened using digital droplet PCR (ddPCR). Populations showing HDR (2 to 10%) were single cell sorted into 96 well plates using FACS and were expanded into clonal cell lines. DNA was extracted from expanded clonal cell populations by establishing a “mirror plate”. Extracted DNA was used directly for genotyping by TaqMan™ qPCR or ddPCR and confirmed by Sanger sequencing. (c) Representative transmitted light microscopy picture of transfected U-CH1 showing satisfactory viability and morphology. (d) Dot plots showing the gating strategy for sorting U-CH1 cells based on TOPRO3 and ATTO-550 fluorescence by FACS.

### 3.7. Screening of transfected bulk populations before isolation of U-CH1 clonal lines

After transfection we screened the bulk population to ensure HDR was present before proceeding to single cell sorting (Figure 3b). For this we utilised digital droplet PCR (ddPCR) as a simpler, faster alternative to MiSeq™ which allowed us to maintain the cells in culture during screening rather than freezing the cells as was done in the iPSC experiments. The partitioning technology of ddPCR enables the detection of rare knock-in events at a frequency as low as 0.1% [36,37]. We designed a ddPCR assay that amplified a small stretch of DNA (~130 bp) surrounding the G177D variant in *TBXT* with fluorescent probes complementary to unedited and edited alleles (Supplementary Figure 4). Whereas NHEJ repair would be expected to disrupt the binding of the primers and probes, producing droplets without fluorescence, alleles repaired by HDR allow amplification of the probes, causing fluorescence that can be quantified. Screening of the 10 transfected populations by ddPCR showed an editing efficiency of 2 to 10%, all of which were taken forward for single cell sorting.

The transfected cells tolerated single cell sorting by FACS as early as 24 to 36 hours after transfection (Figure 3c-d). The single cells required 3 to 4 weeks to expand sufficiently to allow subculture into a “mirror” plate from which genomic DNA was extracted for genotyping. The Lucigen one-step DNA extraction protocol was utilised, allowing direct input of the DNA solution into TaqMan™ qPCR or further ddPCR for high throughput genotyping. Wells showing edited alleles were further characterised by Sanger sequencing to identify off-target alterations.

### 3.8. High throughput screening of hundreds of U-CH1 clones

We screened ~500 transfected clonal lines and isolated seven lines of interest which were free of indels: four wild type, two heterozygous knock-ins and one homozygous knock-in (Figure 4a-c), giving an overall HDR efficiency of 0.6%. In contrast, we identified more than 250 clones in which there was a combined repair process involving both the knock-in and indels introduced by NHEJ (Figure 4b).

**Figure 4.**
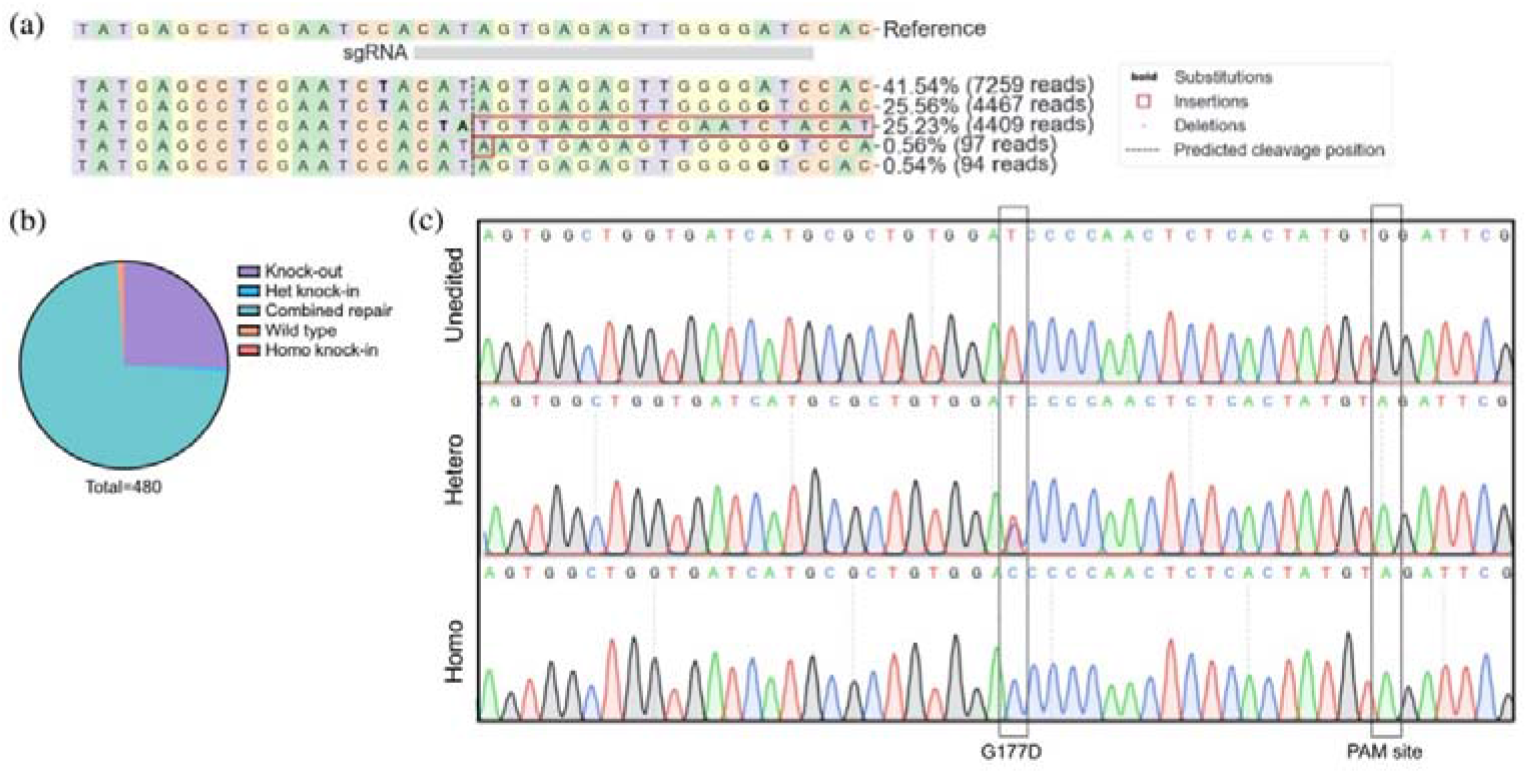
Editing outcomes of U-CH1 cell line. (a) MiSeq™ data from a clonal line, processed by CRISPResso2 [38] showing a greater proportion of reads with knock-in of the blocking mutation (bold A) than G177D SNV (bold G). (b) Breakdown of editing outcomes across ~500 clones expanded from single cells. (c) Sanger sequencing traces of edited clones showing accurate editing of the G177D SNV and blocking mutation.

The relationship between HDR efficiency and cut-to-mutation distance [4,5] was again demonstrated in our experiments: the silent blocking mutation introduced into the ssODN was successfully edited more frequently than the G177D SNV and was edited almost exclusively in a homozygous fashion, in contrast to the G177D SNV which was only edited in one allele (Figure 4c).

Finally, we performed quality assurance on the isolated clones by sequencing 1,000 bp around the site of the edit by Sanger sequencing: we confirmed the presence of the introduced edit and ensured no off-target alterations were present (Supplementary Figure 5).

In summary, we show that editing stem cells and immortalised cell lines at disease-relevant genomic loci is challenging. However, even when constrained by gRNA options, it is possible to boost HDR rates in iPSC and screen the hundreds of potential clones using high throughput technologies. We propose a flowchart which could be used to guide the planning of CRISPR/Cas9 experiments to edit single nucleotide variants (Figure 5).

**Figure 5.**
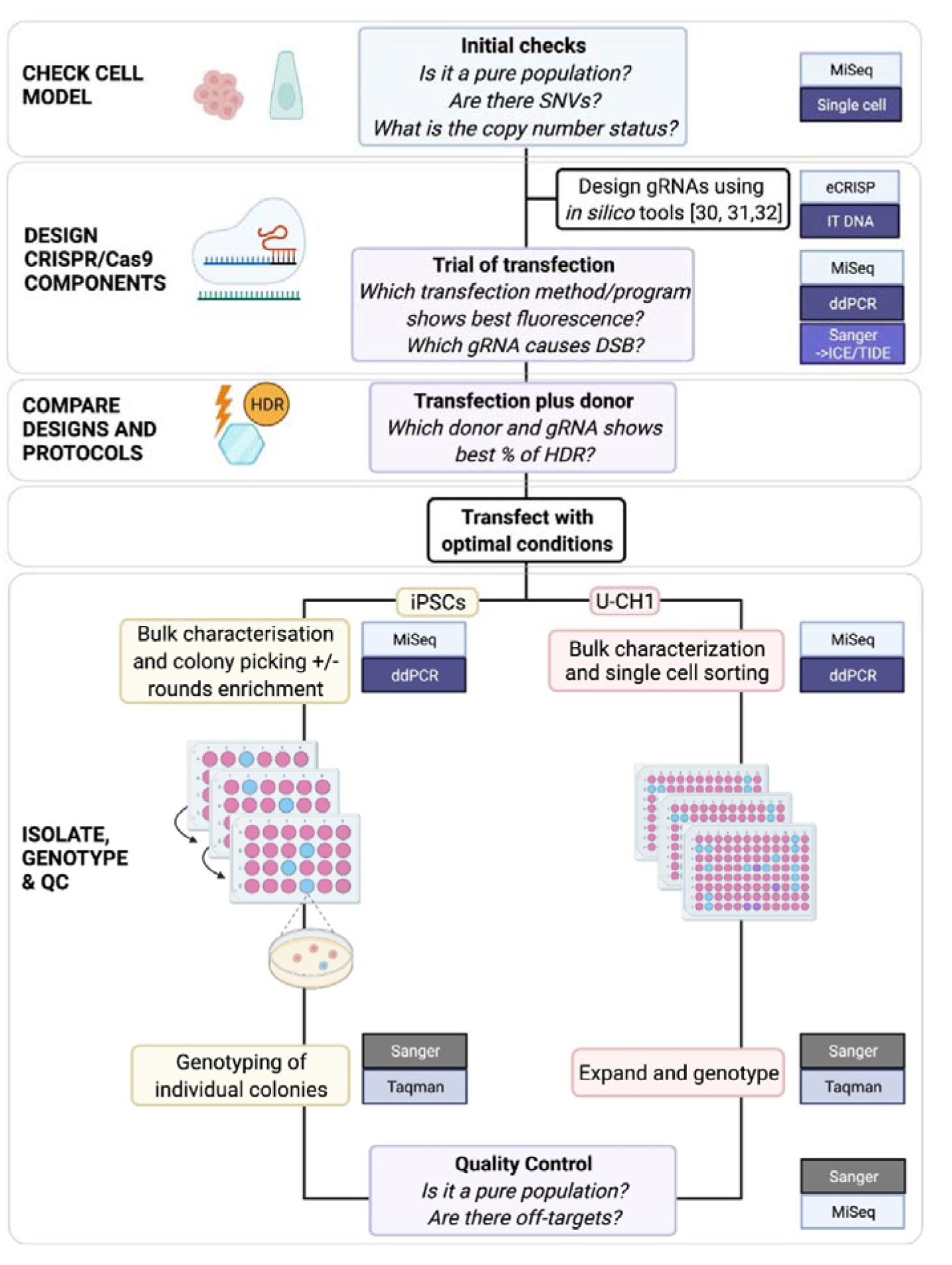
Proposed comprehensive workflow for editing stem cells or immortalised cell lines using CRISPR/Cas9.

## 4. Discussion

The relative ease of designing a gRNA and the high DNA cleavage efficiency of Cas9 make CRISPR/Cas9 an attractive genome editing system. However, the range of parameters that can be modified introduces complexity into experimental design. Furthermore, the low efficiency of HDR necessitates the screening of large numbers of clonal lines to isolate those that have been accurately edited.

Based on our experience, we present an adaptable workflow for improving single nucleotide editing in stem cells and immortalised cell lines using CRISPR/Cas9. For stem cells, the protocol incorporates MiSeq™ for characterising editing outcomes in the transfected bulk population before isolating clonal lines and is suited to cell models for which it is impractical to maintain large numbers of clones. For immortalised cell lines, a rapid screening step is employed after which we proceed to early single cell sorting followed by high throughput genotyping; a protocol that is suited to models for which several clonal lines can be maintained until genotyping is possible.

In our CRISPR/Cas9 experiments, we found that the efficiency of single nucleotide editing varied between transfections. Our rate of HDR at the level of the bulk population was lower [5,7,16] or similar [39] to that of previously published optimisation reports in which bulk population data were analysed. The lower efficiency in our iPSC experiments can be explained by the suboptimal cut-to-mutation distance which is determined by the location of the gRNA. For both models, the computational design tools suggested four to six gRNAs, but we found that only one gRNA showed efficacy *in vitro* in each model, thus highlighting the need to test *in silico* predictions [40]. However, when the gRNA or cut-to-mutation distance cannot be altered, we show that combining an asymmetric donor template, cold shock and HDR enhancer improves the efficiency of accurate repair, corroborating the findings of others [7,16].

Once editing protocols have been optimised, the next challenge is identifying the subset of rare edited cells within a transfected population and isolating these as viable clonal lines. Sanger sequencing, the T7 endonuclease 1 mismatch detection assay [41] and TaqMan™ genotyping are not sufficiently sensitive to screen transfected bulk populations for cells that have undergone single nucleotide editing [36]. Computational modelling tools such as the Synthego ICE (Inference of CRISPR Edits) [42] and Tracking Indels by Decomposition (TIDE) tools [43] can be employed to analyse Sanger sequencing generated from bulk cell populations to infer the composition of editing outcomes [42,43], however their sensitivity is low for rare events. We therefore chose to screen transfected populations using two quantitative methods that can detect HDR rates as low as <0.1%: MiSeq™, although this is costly and requires cells to be frozen while waiting for results, and ddPCR [36,37] which can be performed rapidly in-house while cells were maintained in culture.

Most previously published protocols report HDR efficiency at the level of bulk populations and do not provide details of HDR efficiency following the isolation of clonal lines. The isolation of viable clonal lines, achieved through cell sorting or colony picking, is required for functional experiments. Intuitively, we thought that the efficiency would be similar to that of the bulk populations but found that this was not the case. Despite promising rates of HDR at the bulk level (12%) these converted poorly to the isolation of clonal lines giving an overall HDR accurate efficiency of 1%. The reasons for this are not clear but it is possible that editing genes that are critical to the biology of our cell models may have caused a selective disadvantage for edited clones, resulting in the isolation of fewer than expected clonal lines. It is also not possible to screen all clones in the bulk population and the random sampling of a small subset for single cell isolation may not be representative of the bulk.

We also highlight that once clonal lines are obtained, important quality control steps need to be undertaken to ensure purity and to exclude off-target alterations. To this end, we employed MiSeq™ which offers high depth analysis of around 200bp region around the cut site, ensuring scarless editing. Assessment of copy number at the locus of interest is also recommended to ensure MiSeq™ data are not confounded by deletions [43]. Moreover, we show that Sanger sequencing a larger region around the variant of interest should be considered to exclude off-targets alterations at distances of 1-2 Kb from the cut site.

Besides CRISPR/Cas9, the more recently described base and prime editing systems offer alternative methods for introducing small substitutions without generating a DSB and relying on cellular repair mechanisms [45]. As with CRISPR methods, these systems are still constrained by sequence requirements: base editing is unsuitable for editing a single nucleotide within repeats (bystander effect) [46] and classical prime editors have the same PAM requirement [25,45]. Besides the Cas9 nuclease, several other bacterial nucleases are available, and these may offer alternatives when genomic region constraints make the use of Cas9 challenging. The smaller size of other nucleases may for example allow greater accessibility to chromatin [47]. The adaptable workflow that we describe, including screening methods and quality control steps, is applicable to these alternative editing systems.

## 5. Conclusions

Elucidating the biological impact of genetic variants plays an important role in understanding disease and has clinical implications. CRISPR/Cas9 is a powerful tool for modelling these variants, however successful editing of single nucleotides is variable and is influenced by the genomic context of the variant and the cellular model that is employed. Investment in such experiments can be time-consuming and costly. Here we provide a step-wise approach that helps to manage and mitigate some of the limitations of the CRISPR/Cas9 editing system. This workflow can be used to generate isogenic lines that allow the functional study of pathogenic and likely-pathogenic single nucleotide variants which are associated with cancer.

## Author Contributions

Conceptualization, I.U., S.A., G.L.B., A.M.F, L.C.; methodology, I.U., S.A., A.M.F. and L.C.; formal analysis, I.U., E.H. and C.C; investigation, I.U., L.L. and L.C.; writing— original draft preparation, I.U. and L.C.; writing—review and editing, L.L., S.A., E.H., C.C., A.M.F, L.C..; visualization, I.U. and L.C.; supervision, L.C. and A.M.F.; funding acquisition, I.U., G.L.B., A.M.F and L.C.. All authors have read and agreed to the published version of the manuscript.

## Funding

This work was funded by the Tom Prince Trust (A.M.F.), The Pathological Society of Great Britain and Ireland (grant number TSGS041903) (I.U. and A.M.F), Cancer Research UK (University College London (CRUK UCL) Centre Award C416/A25145) (A.M.F, L.C. and G.B.), the Royal National Orthopaedic Hospital NHS Trust R&D Department, Bone Cancer Research Trust, Chordoma Foundation. A.M.F. was supported by the National Institute for Health Research, the University College London Hospitals Biomedical Research Centre, and the Cancer Research UK University College London Experimental Cancer Medicine Centre. I.U. was supported by Chordoma UK. S.A. was supported by the Hardy Keinan fellowship.

## Acknowledgments

We acknowledge Sunniyat Rahman, Samuel Weeks, Daniel Wetterskog, Yang Li, Iben Lyskjaer and Florian Merkle for their invaluable advice in setting up our protocols.

## Conflicts of Interest

The authors declare no conflict of interest.

## Supplementary data

**Supplementary Table 1.** STR (Short Tandem Repeat) analysis results for U-CH1, used in the study.

| Marker | U-CH1 profiles |  | Database profiles |  |
| --- | --- | --- | --- | --- |
|  | Allele 1 | Allele 2 | Allele 1 | Allele 2 |
| AMEL | X | Y | X | Y |
| CSF1PO | 10 | 11 | 10 | 11 |
| D13S317 | 11 | 13 | 11 | 13 |
| D16S539 | 12 | 13 | 12 | 13 |
| D18S51 | 15 | 15 | 15 | 15 |
| D21S11 | 28 | 29 | 28 | 29 |
| D3S1358 | 15 | 15 | 15 | 15 |
| D5S818 | 11 | 12 | 11 | 12 |
| D7S820 | 9 | 12 | 9 | 12 |
| D8S1179 | 10 | 15 | 10 | 15 |
| FGA | 20 | 21 | 20 | 21 |
| Penta D | 11 | 11 | 11 | 11 |
| Penta E | 7 | 10 | 7 | 10 |
| TH01 | 7 | 7 | 7 | 7 |
| TPOX | 8 | 11 | 8 | 11 |
| vWA | 17 | 17 | 17 | 17 |
| AMEL | X | X | X | X |

**Supplementary Table 2.**
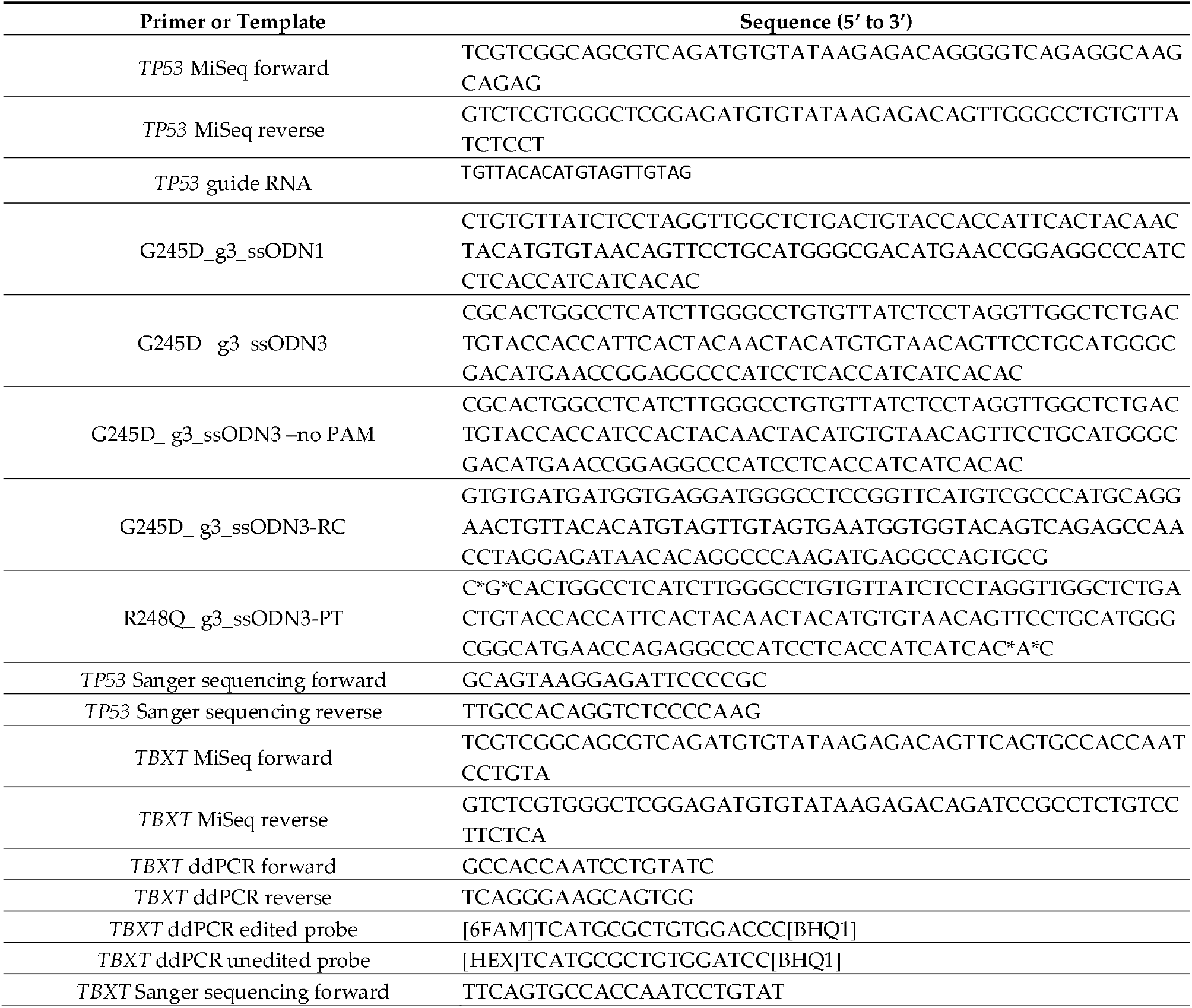
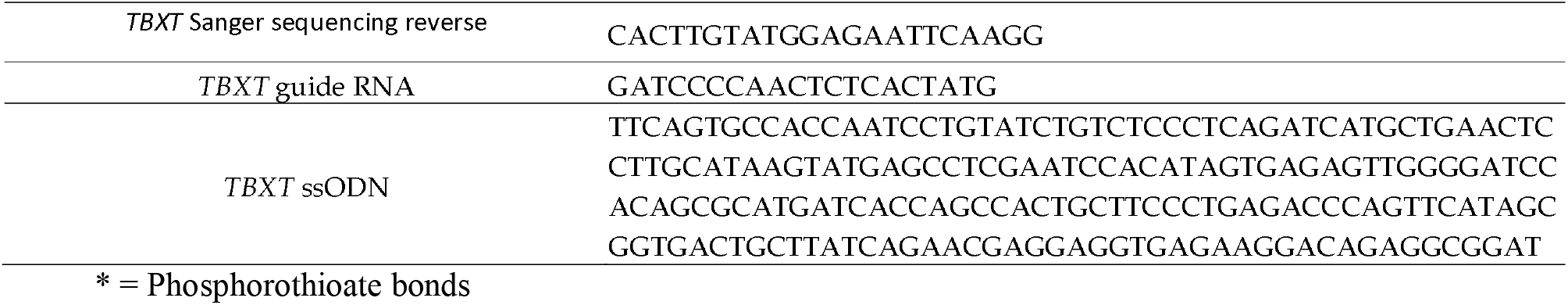
List of primers, guides, donors used in this study.

**Supplementary Figure 1.**
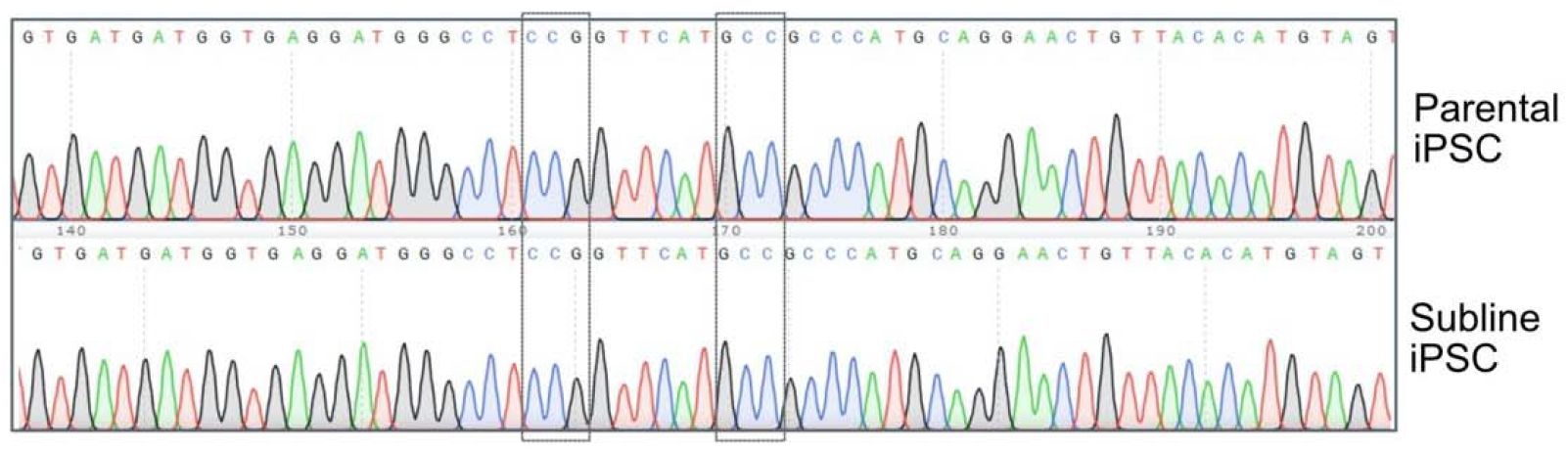
Sanger sequencing of TP53 from colony-picked iPSCs. Representative Sanger sequencing traces of TP53 exon 7 for one of the five tested sublines: all sublines were free of SNVs in the TP53 sequence surrounding the G245D and R248Q loci (highlighted by boxes) and were wild type for the G245D and R248Q SNVs.

**Supplementary Figure 2.**
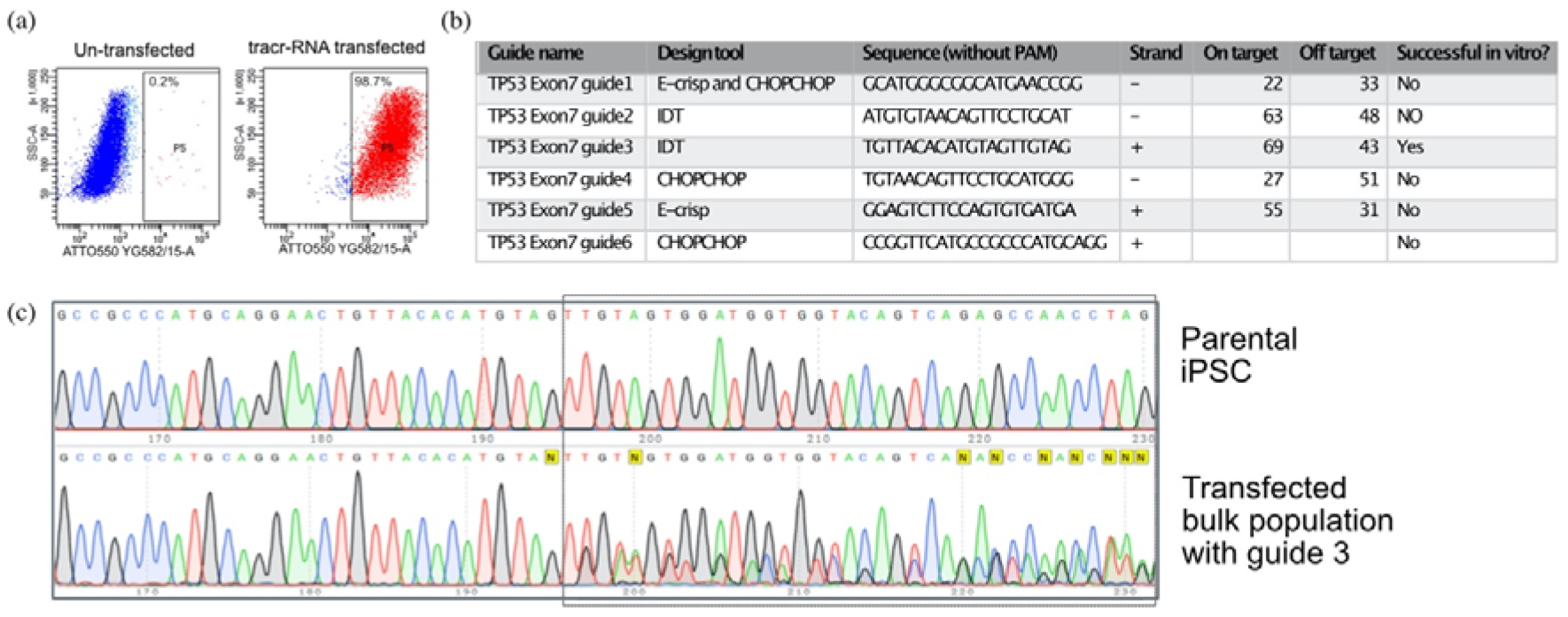
iPSC after transfection with RNP containing *TP53* exon 7 guide RNA 3. (a) Dot plots showing 98.7% ATTO-550-positive cells by FACS after transfection with the gRNA-tracrRNA duplex. (b) Sequence and information on all designed and tested gRNAs. (c) Sanger sequencing traces of the parental iPSC showing the reference sequence (top) and the bulk population transfected with the successful gRNA (*TP53* exon 7 guide 3), showing evidence of repair by NHEJ at the predicted cut site (bottom).

**Supplementary Figure 3.**
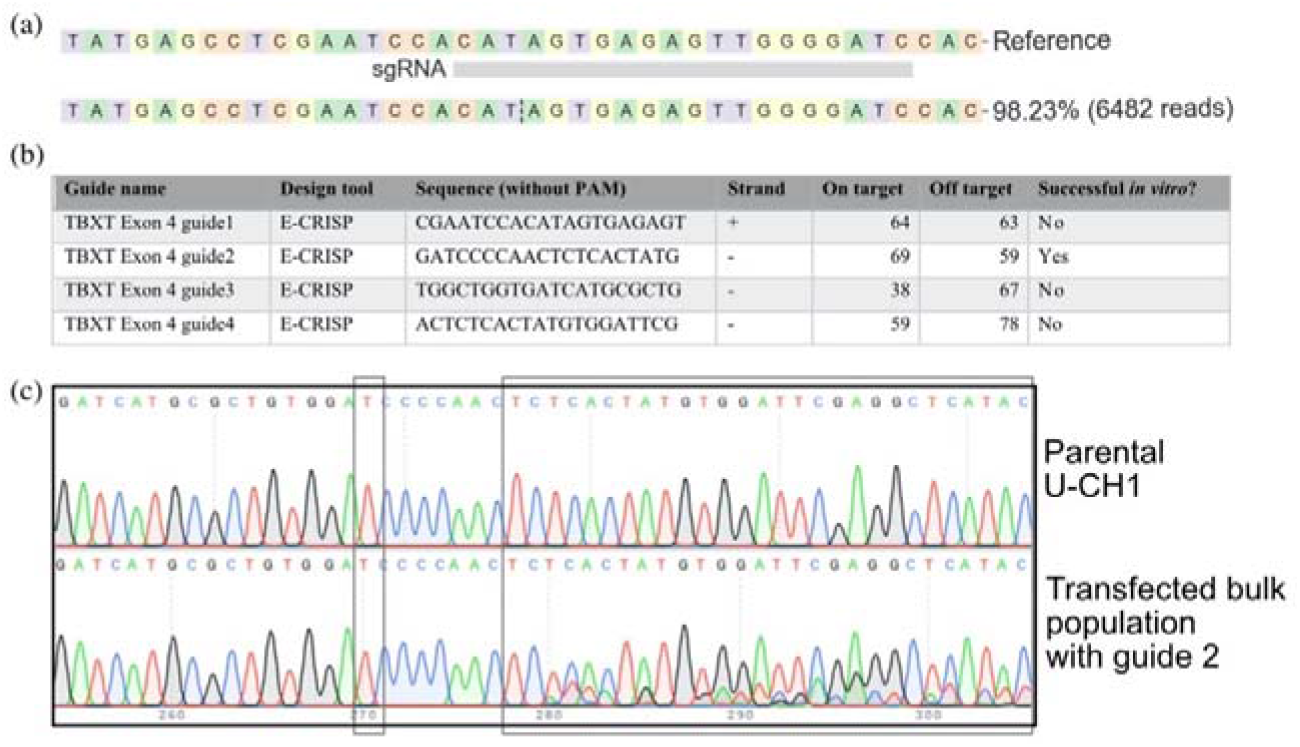
CRISPR/Cas9 gRNA design for U-CH1 chordoma cell line. (a) MiSeq™ results from the U-CH1 parental cell line analysed by CRISPResso2 [38] showing a genetically pure starting population, homozygous for the variant allele at the G177D SNV. (b) Sequence and information on all designed and tested gRNAs. (c) Sanger sequencing traces of the parental U-CH1 cells showing the reference sequence (top) and the bulk population transfected with the successful gRNA, showing evidence of repair by NHEJ at the predicted cut site (bottom). Dashed box on the left highlights the G177D SNV, and on the right highlights the repaired region downstream of the cut site.

**Supplementary Figure 4.**
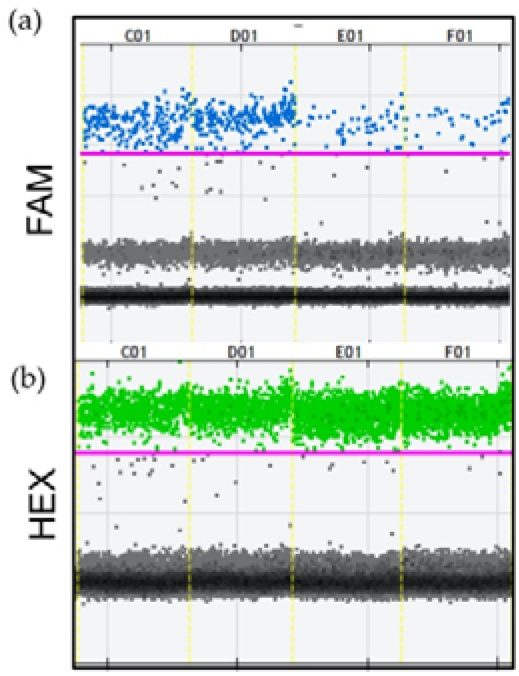
Dot plot of ddPCR assay for edited and unedited alleles in U-CH1. (a) Blue droplets (FAM) represent the edited allele and (b) green droplets (HEX) represent the reference allele.

**Supplementary Figure 5.**
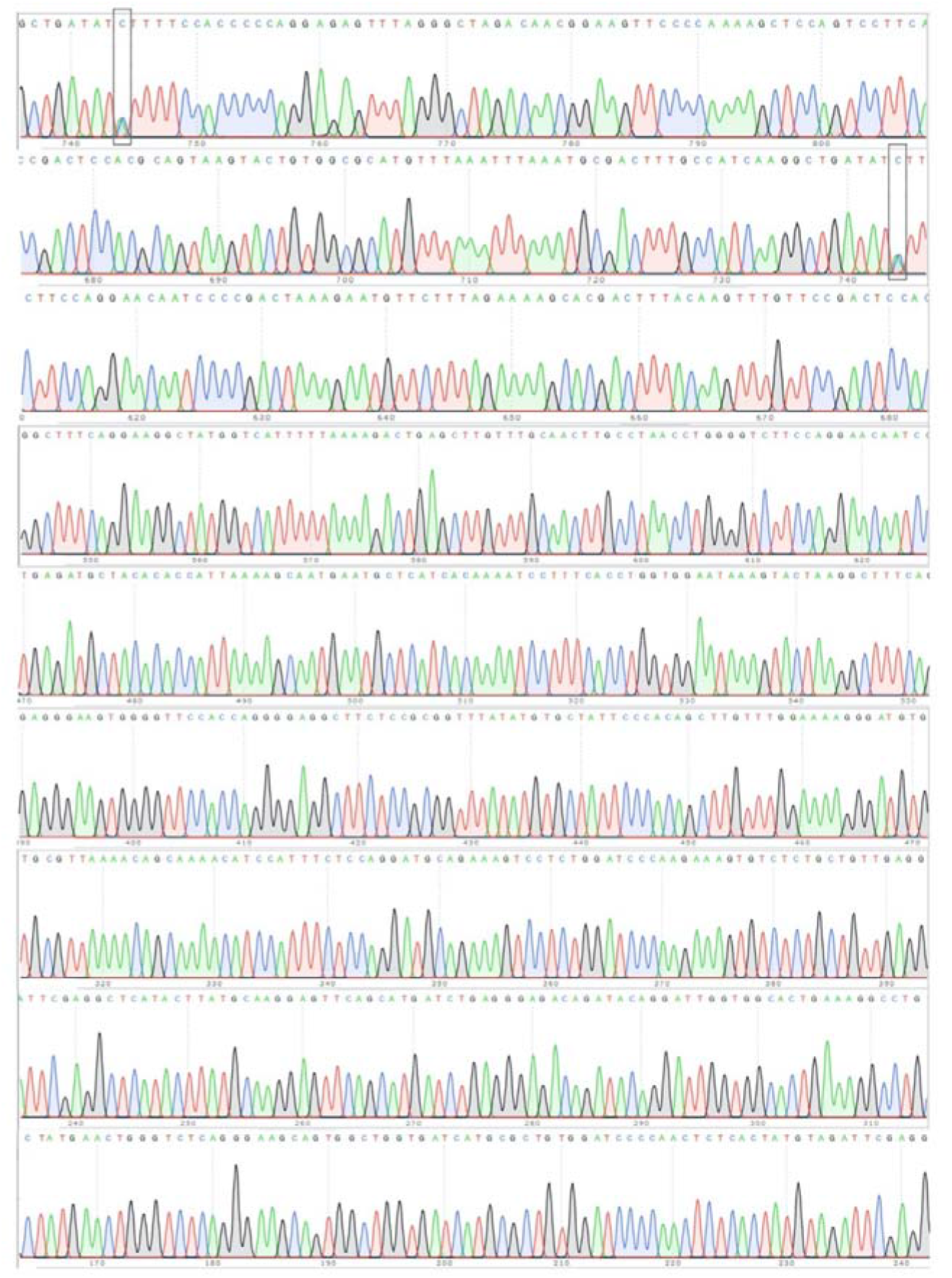
Quality assurance of edited U-CH1 clonal lines. Sanger sequencing traces of 1,000 bp around the site of the edit in one heterozygous edited clone showing accurate editing of the G177D SNV in *TBXT* and no off-target alterations. Dashed boxes indicate the G177D SNV.

## Notes

### Competing Interest Statement

The authors have declared no competing interest.

