## Supplementary figures for "An efficient and adaptable workflow for editing disease-relevant single nucleotide variants using CRISPR/Cas9"

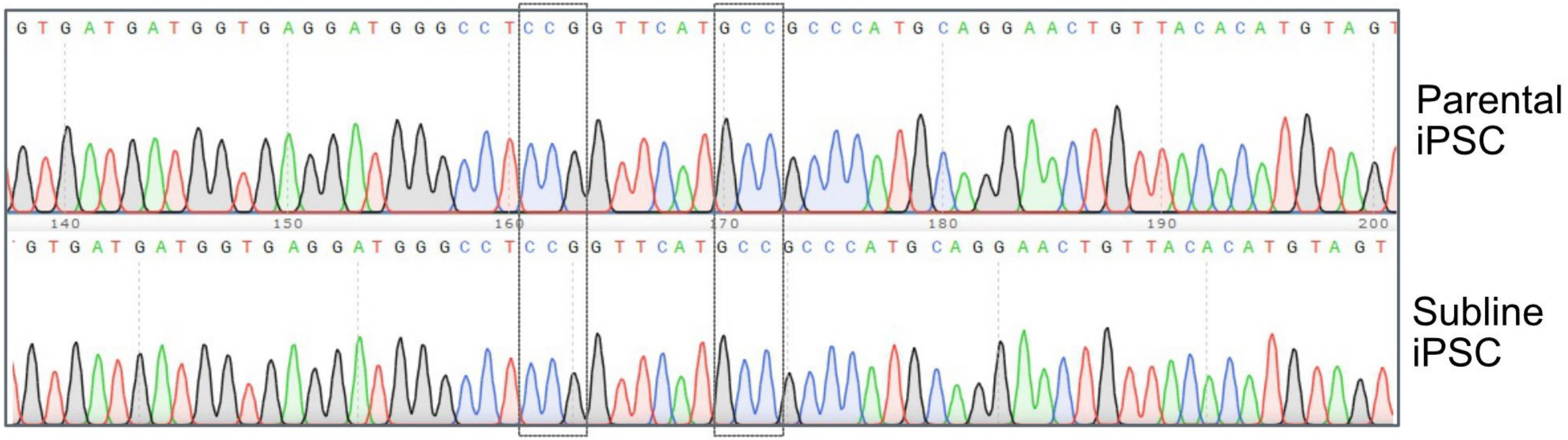

GTGATGATGGTGAGGATGGGCCTCCGGTTCATGCCGCCCATGCAGGAACTGTTACACATGTAGT
140
150
160
170
180
190
200
GTGATGATGGTGAGGATGGGCCTCCGGTTCATGCCGCCCATGCAGGAACTGTTACACATGTAGT
Parental iPSC
Subline iPSC

(a)

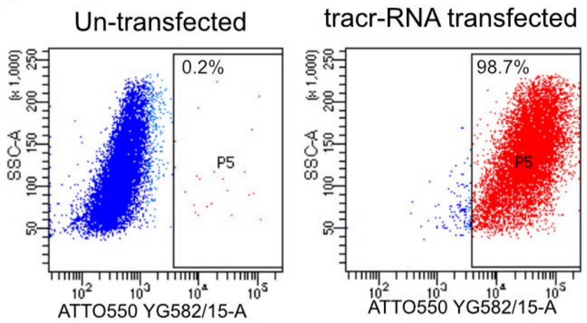


(b)

| Guide name | Design tool | Sequence (without PAM) | Strand | On target | Off target | Successful in vitro? |
|---|---|---|---|---|---|---|
| TP53 Exon7 guide1 | E-crisp and CHOPCHOP | GCATGGGCGGCATGAACCGG | – | 22 | 33 | No |
| TP53 Exon7 guide2 | IDT | ATGTGTAACAGTTCCTGCAT | – | 63 | 48 | NO |
| TP53 Exon7 guide3 | IDT | TGTTACACATGTAGTTGTAG | + | 69 | 43 | Yes |
| TP53 Exon7 guide4 | CHOPCHOP | TGTAACAGTTCCTGCATGGG | – | 27 | 51 | No |
| TP53 Exon7 guide5 | E-crisp | GGAGTCTTCCAGTGTGATGA | + | 55 | 31 | No |
| TP53 Exon7 guide6 | CHOPCHOP | CCGGTTCATGCCGCCCATGCAGG | + | | | No |

(c)

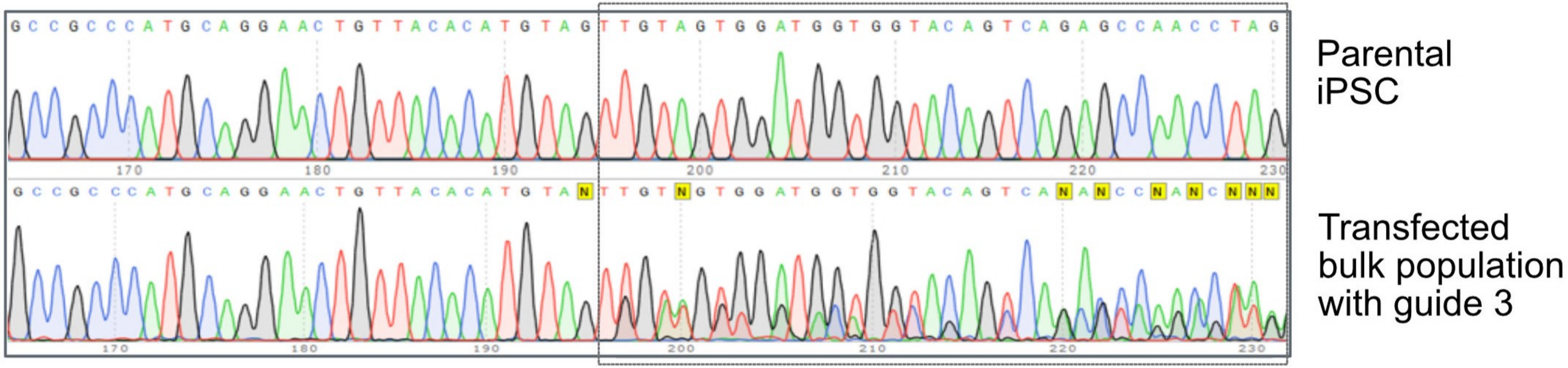

(a)

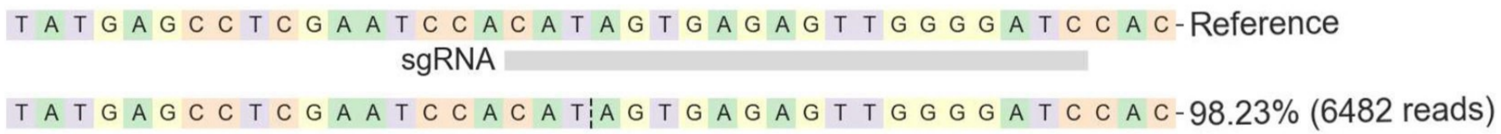


(b)

| Guide name | Design tool | Sequence (without PAM) | Strand | On target | Off target | Successful *in vitro*? |
|---|---|---|---|---|---|---|
| TBXT Exon 4 guide1 | E-CRISP | CGAATCCACATAGTGAGAGT | + | 64 | 63 | No |
| TBXT Exon 4 guide2 | E-CRISP | GATCCCCAACTCTCACTATG | - | 69 | 59 | Yes |
| TBXT Exon 4 guide3 | E-CRISP | TGGCTGGTGATCATGCGCTG | - | 38 | 67 | No |
| TBXT Exon 4 guide4 | E-CRISP | ACTCTCACTATGTGGATTCG | - | 59 | 78 | No |

(c)

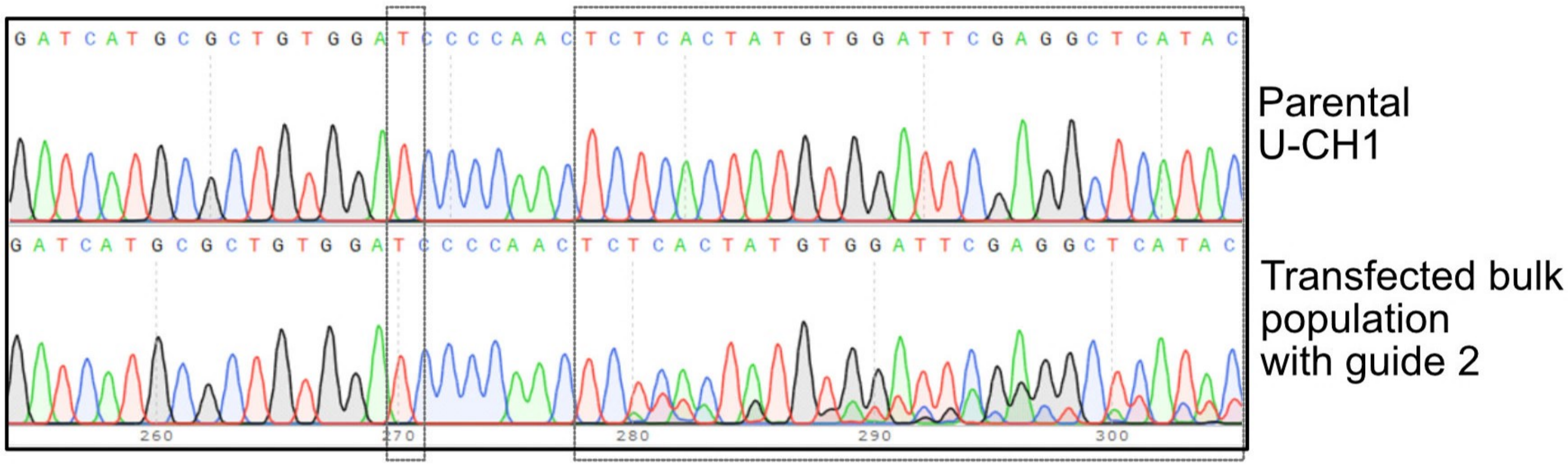

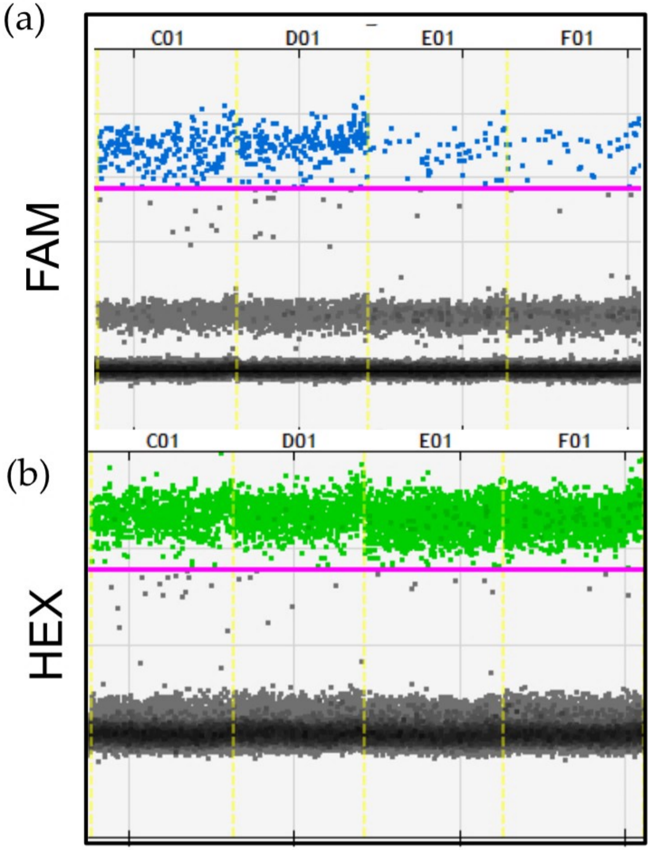
(a)
C01
D01
E01
F01
FAM
(b)
C01
D01
E01
F01
HEX

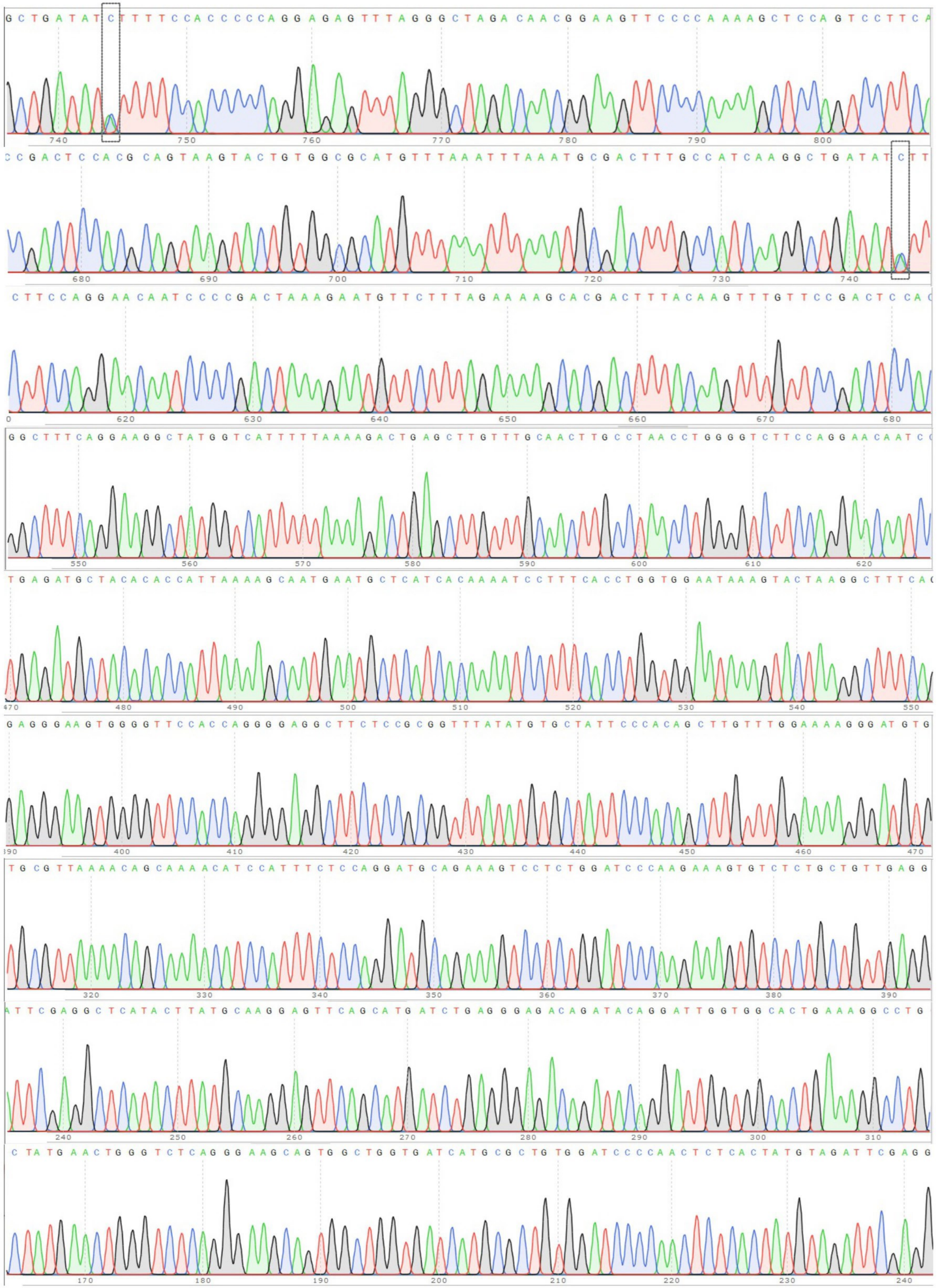
